# MOSTPLAS: A Self-correction Multi-label Learning Model for Plasmid Host Range Prediction

**DOI:** 10.1101/2024.07.31.606102

**Authors:** Wei Zou, Yongxin Ji, Jiaojiao Guan, Yanni Sun

## Abstract

Plasmids play an essential role in horizontal gene transfer among diverse microorganisms, aiding their host bacteria in acquiring beneficial traits like antibiotic and metal resistance. Identifying the host bacteria where a plasmid can transfer, replicate or persist provides insights into how plasmids promote bacterial evolution. Plasmid host range prediction tools can be categorized as alignment-based and learning-based. Alignment-based tools have high precision but fail to align many newly sequenced plasmids with characterized ones in reference databases. In contrast, learning-based tools help predict the host range of these newly discovered plasmids. Although previous researches have demonstrated the existence of broad-host-range (BHR) plasmids, there is no database providing their detailed and complete host labels. Without adequate well-annotated training samples, learning-based tools fail to extract discriminative feature representations and obtain limited performance. To address this problem, we propose a self-correction multi-label learning model called MOSTPLAS. We design a pseudo label learning algorithm and a self-correction asymmetric loss to facilitate the training of multi-label learning model with samples containing some unknown missing positive labels. Experimental results on multi-host plasmids generated from the NCBI RefSeq database, metagenomic data, and real-world plasmid sequences with experimentally determined host range demonstrate the superiority of MOSTPLAS.

## Introduction

Plasmid is a small circular, double-stranded and independently replicated DNA molecule within a cell [1]. It is mostly found in bacteria and physically separated from chromosomal DNA. In some cases, the genes encoded by plasmids provide bacteria with beneficial traits such as antibiotic resistance [2], metal resistance [3], and virulence catabolic capacity [4]. With these genetic advantages, plasmids gain widespread attention among researchers [5, 6]. To obtain comprehensive understanding of how plasmids facilitate the evolution of bacteria, we first need to explore the relations between plasmids and their host bacteria.

The host range of a plasmid refers to the microorganisms where a plasmid can transfer, replicate, or persist [7]. Accordingly, plasmids can be classified into narrow-host-range (NHR) plasmids and broad-host-range (BHR) plasmids. BHR plasmids refer to a group of plasmids that can replicate and stably maintain among organisms of different phylogenetic groups [7].

BHR plasmids are closely associated with horizontal gene transfer among bacteria of different taxonomic groups. Correctly predicting the host range of BHR plasmids is beneficial for us to explore how plasmids devote to the spread of antimicrobial resistance genes among human pathogens [8]. In addition, with discovery of more BHR plasmids, their replicons can be used to create recombinant vectors that can replicate and express proteins in different bacteria [9]. These vectors can further sever as a potential medical treatment in gene therapy.

Existing tools designed for plasmid host range prediction can be summarized as alignment-based tools [10, 11] and learning-based tools [12, 13]. The assumption behind alignment-based plasmid host range prediction tools is that plasmids with high sequence similarity tend to share the same host range. Learning-based plasmid host range prediction tools first extract and aggregate various kinds of sequence features of plasmids, such as *k*-mer frequency, GC content, codon usage information, replicon type, mobilization (MOB) type, etc. Then, these features are fed into many kinds of learning models such as convolutional neural network, transformer, random forest to predict the host range of input plasmid sequence. Compared with alignment-based tools, learning-based tools have higher sensitivity for predicting the host range of novel plasmids that may have no alignments with characterized plasmids.

Although learning-based plasmid host range prediction tools have superiority on detecting the host range of newly discovered plasmids, there are still some unsolved limitations. Previous researches and experiments [14, 15, 16] have demonstrated the existence of BHR plasmid that have multi-host range labels among different phylogenetic groups. However, there are no available databases providing detailed and complete host ranges of BHR plasmids. The complete RefSeq plasmid sequences downloaded from NCBI database are solely annotated with the host label from which they are isolated. Without sufficient well-annotated training samples, it is difficult for learning-based tool to realize the potential of deep learning models on conducting comprehensive host prediction for plasmids that may have multiple host labels. Another challenge lies in the highly imbalanced host labels associated with each plasmid, where the number of non-host bacteria can significantly exceed the number of actual host organisms. And thus, negative labels dominate the model parameter optimization process, leading to undesirable performance. To fully leverage the powerful learning models nowadays, plasmid host range prediction can be formulated as a multi-label learning problem with missing labels.

In the field of computer vision, many algorithms have been proposed and achieved the state-of-the-art performance on the problems of multi-label learning with missing labels. Among them, Lin et al. proposed Focal Loss [17] to cope with the problem that most locations in a sample are easy negatives that contribute no useful learning information for model training in objection detection task. Focal Loss employed a hyper-parameter to differentiate the importance of hard and easy examples, so that those easily classified samples were down-weighted. On the basis of Focal Loss, Tal et al. proposed Asymmetric Loss [18], which discarded the loss of negative samples with very low predicted probabilities and explicitly adopted different focus parameters for positive and negative classes to control their contributions tn model training. For the task of facial expression recognition, the annotations of training samples involve uncertainty caused by ambiguous facial expressions, low-quality facial images, and the subjectiveness of annotators. To mitigate the influence of potential false negative labels, Wang et al. [19] proposed a relabeling mechanism that replaced the original label of an input with a new pseudo label if the maximum prediction probability is higher than the corresponding probability of the given label by a significant margin.

Inspired by these algorithms, we propose a self-correction multi-label learning model named MOSTPLAS for genus-level plasmid host range prediction. In MOSTPLAS, we design a pseudo label generation algorithm to assign some reliable labels for each plasmid sequence, which provides extra supervision information for the training of multi-label learning model. We also replace commonly used binary cross-entropy loss with our self-correction asymmetric loss to reduce the negative influence of extreme imbalance between the number of positive and negative labels. With the aforementioned modification, MOSTPLAS can perform well under the challenging scenario that the training samples are only annotated with one positive label.

The main contributions of our paper are summarized below:

1. To the best of our knowledge, this is the first attempt to implement a multi-label learning model for genus-level plasmid host range prediction.
2. We design a pseudo label generation algorithm to cope with the problem of lacking complete host label annotations for the training of multi-label learning model. To alleviate the influence of large imbalance between the number of positive labels and negative labels in RefSeq plasmid sequences, we also implement a self-correction asymmetric loss to help the model extract discriminative feature representation for each class.
3. To demonstrative the superiority of our proposed MOSTPLAS, we perform a series of experiments on multi-host plasmid test set generated from the NCBI RefSeq database, metagenomic data and real-world plasmid sequences with experimentally determined host range. Experiment results show that our model performs better than both alignment-based tools and existing learning-based tools.

## Material and method

### Overview of our self-correction multi-label learning model

With an input complete plasmid sequence, MOSTPLAS aims at predicting all the possible host labels at genus level in an end-to-end manner. The framework is sketched in Fig. 1.

**Fig. 1.**
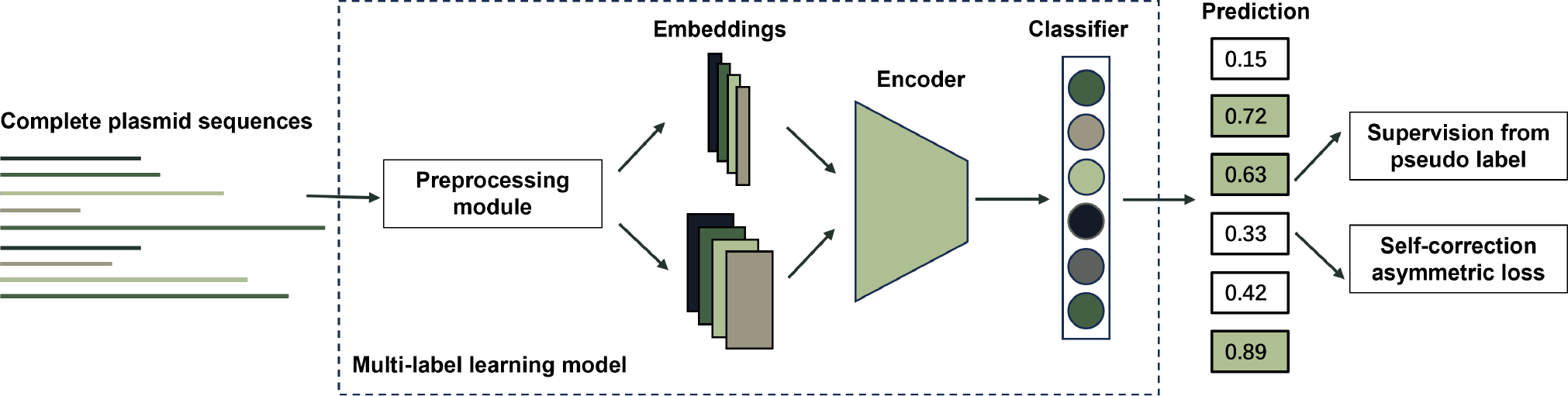
Framework of our proposed MOSTPLAS. Given a complete plasmid sequence, the preprocessing module first converts it into an embedding with fixed length. Based on different designs of encoder in our MOSTPLAS, the embedding can be a vector or a matrix. The embedding is then fed into the encoder to extract discriminative feature representation for classification. At last, the classifier outputs the predicted possibility of input plasmid sequence corresponding to different genera. In order to improve the performance of multi-label learning models, we adopt pseudo labels to provide extra supervision information for parameter updating and replace traditional binary cross-entropy loss with our proposed self-correction asymmetric loss.

As the complete plasmid sequences are of different lengths, in MOSTPLAS, we incorporate a preprocessing module to convert the input sequences into embeddings with fixed size in order to meet with the model input requirement. The embeddings are then sent into the encoder to extract discriminative feature representations. We design three kinds of encoders: Neural Network (NN), Convolutional Neural Network (CNN), and a multi-model integration framework (NN+CNN). The detailed configuration is presented in the supplementary file. The feature representations extracted by the encoder are further passed through the classifier to predict the possibilities that the input plasmid sequences belonging to different genera. In MOSTPLAS, the classifier is implemented with a fully-connected layers with the same amount of hidden neurons as the number of total genera.

To cope with the limitation that the RefSeq plasmid sequences downloaded from NCBI database only contain single host label, we design a pseudo label generation algorithm to perform data augmentation on the annotations of training samples. These reliable pseudo labels can be regarded as extra supervision information for the training of multi-label learning model. We also design a self-correction asymmetric loss to mitigate the preference of traditional binary cross-entropy loss that pays more attention to the negative labels during model parameter optimization. In addition, with the help of self-correction asymmetric loss, the model itself can recognize some potential host labels during model training. In the following sections, we will elaborate the algorithm details.

### Pseudo label generation

Lacking sufficient well-annotated samples, it is difficult for a deep learning model to take full advantage of its ability to learn discriminative features for downstream classification or regression tasks [20]. However, under many scenarios, labeling a large dataset is very expensive and even not feasible if the annotation of samples requires expert knowledge or extensive experiments. In semi-supervised learning, many pseudo labelling algorithms [21, 22, 23] have been proposed and demonstrated their effectiveness on leveraging partially labeled or unlabeled samples during model training to help recent deep learning models achieve remarkable gains in performance.

For our task of plasmid host range prediction, there are no available databases providing the complete host range annotations of plasmids that can replicate in multiple hosts. The incomplete training data prevents deep learning models from extracting compact feature representations that contains the characteristics of all the hosts for BHR plasmids, which further limits the performance of learning-based tool on recognizing all the host labels of BHR plasmids. To cope with this challenge, we borrow the idea from recent pseudo labelling algorithms.

Previous research concludes that plasmids with high sequence similarity tend to share the same host range [24]. Experiments also find that some plasmids have multiple origins of replication, and these origins are activated separately in different types of hosts [25]. These characteristics suggest that we can design a pseudo label generation algorithm based on the distribution of plasmid-encoded proteins among different genera. As the host adaptation is largely determined by the encoded proteins in plasmids, our goal is to identify potential extra hosts based on the protein similarity and organization. With our pseudo label generation algorithm, some plasmids will obtain an extra host label as output. The flowchart of our pseudo label generation algorithm is illustrated in Fig. 2.

**Fig. 2.**
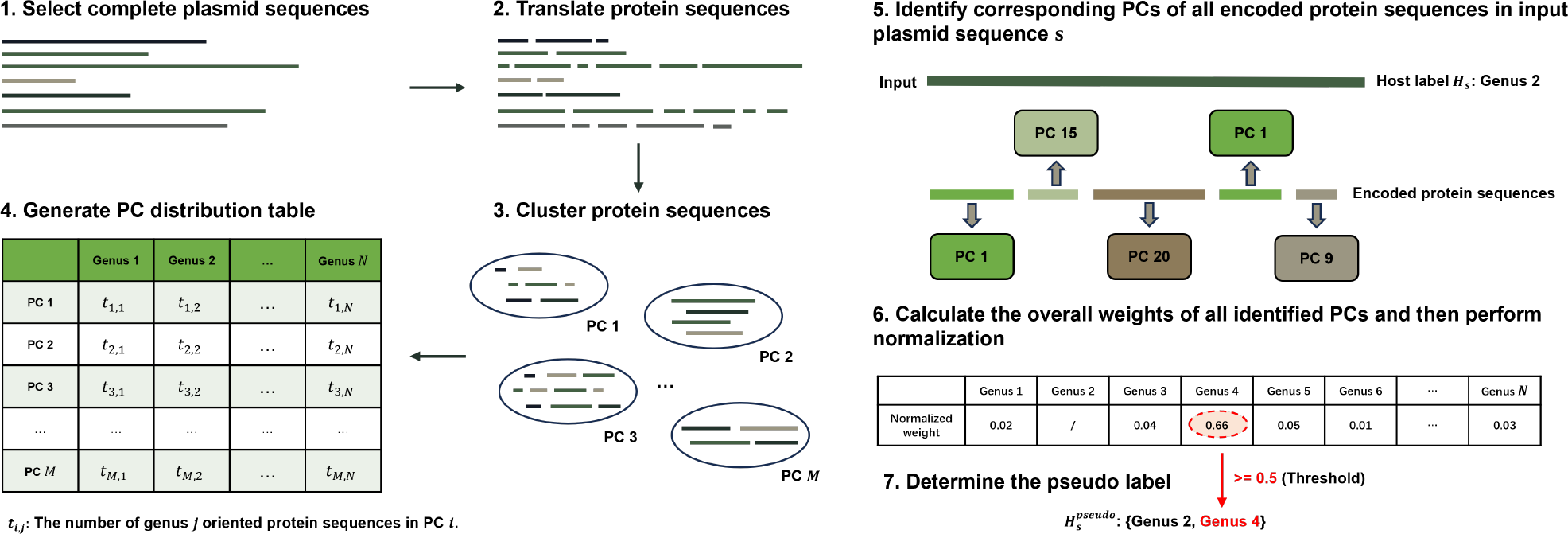
Flowchart of our pseudo label generation algorithm. To obtain reliable pseudo labels, We downloaded and selected the complete plasmid sequences from the NCBI RefSeq database. Then, we used Prodigal to translate the protein sequences and employed CD-HIT to cluster all the protein sequences. The clustered proteins (i.e. protein families/clusters) and their distributions along plasmids of different genera are used to compute the significance score of a protein family to a genus. With the significance score, we first identify the corresponding PCs of all encoded protein sequences in an input plasmid sequence. Afterwards, we averaged the significance scores of each identified PC and perform normalization on the overall weights of all PCs in input plasmid. If the normalized weight of one genus is larger than the threshold, we determine it as the pseudo label.

We generate the pseudo label for selected complete plasmid sequences in the training set. We first downloaded the plasmid sequences from the NCBI RefSeq database. The latest version is released on 9 March 2024 and contains 54,794 sequences. We selected the sequences with the assembly level as complete genome and downloaded their corresponding host lineages. As the focus of this work is the bacterial hosts, we removed the sequences whose hosts do not belong to bacteria. We also discarded the sequences corresponding to genus that contains less than 10 sequences as the number of samples is not sufficient for model training. As a result, we obtained 41,074 complete plasmid sequences corresponding to 216 genera in total.

To obtain the protein organization, we adopted Prodigal [26] for gene prediction and translation in all the selected complete plasmid sequences. Then, we applied CD-HIT [27] (with threshold=0.9) to cluster all the protein sequences. We summarized the number of protein sequences corresponding to different genera within each protein cluster (PC) and obtained a protein cluster distribution table shown in Fig. 2. We will use this table to compute the importance of a PC to a host genus. Then, we will derive the extra pseudo label of a plasmid sequence by the corresponding importance of its encoded proteins.

In information retrieval, term frequency–inverse document frequency (TF-IDF) is a commonly used measurement of the importance of a word to a document in a collection or corpus [28].

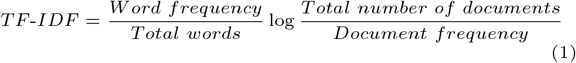

We can also adopt TF-IDF to evaluate the significance of one PC to a genus, i.e., the PCs generated by CD-HIT can be regarded as the words, and all the genera can be considered as different types of documents. And thus, if a PC appears more frequently than other PCs in this genus, this PC has a higher significance to this genus. On the contrary, if this PC contains protein sequences of all the genera in the same time, this PC has a low TF-IDF score.

However, there are still some limitations on applying TF-IDF to calculate the significance of one PC to a genus. If a set of homologous protein sequences in one PC appear in all genera, the TF-IDF score of this PC is zero. In our CD-HIT clustering results, there are a large number of PCs that include protein sequences of all the genera, which results in no difference in their significance. Essentially, TF-IDF neglects the appearance frequency of a PC among different genera. As a result, it may not accurately reflect the importance of a PC to a genus. In our application, a PC *i* is importance to a genus *j* if this PC *i* appears more frequently than other PCs in genus *j* and meanwhile appears more frequently in genus *j* than other genera. In order to incorporate the normalized frequency of a PC in one genus and its relative frequency across different genera, we design a TF-IDF^*pro*^ to evaluate the significance of a protein cluster *i* to a genus *j*:

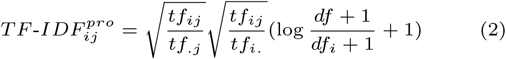

where *tf*_*ij*_ is the number of genus *j* oriented protein sequences in PC *i, tf*_.*j*_ = Σ_*i*_ *tf*_*ij*_, *tf*_*i*._ = Σ_*j*_ *tf*_*ij*_, *df* is the number of total genera in our selected complete plasmid sequences, *df*_*i*_ is the number of genera involves in PC *i*.

To visually show the difference between TF-IDF and TF-IDF^*pro*^ significance scores, we first sorted the PCs based on the total number of protein sequences in each PC. Then, we used heat maps to present the TF-IDF significance scores and TF-IDF^*pro*^ significance scores of the top and last 50 PCs in Fig. 3. In each heat map, darker color represents higher significance score of a protein cluster *i* to a genus *j*.

**Fig. 3.**
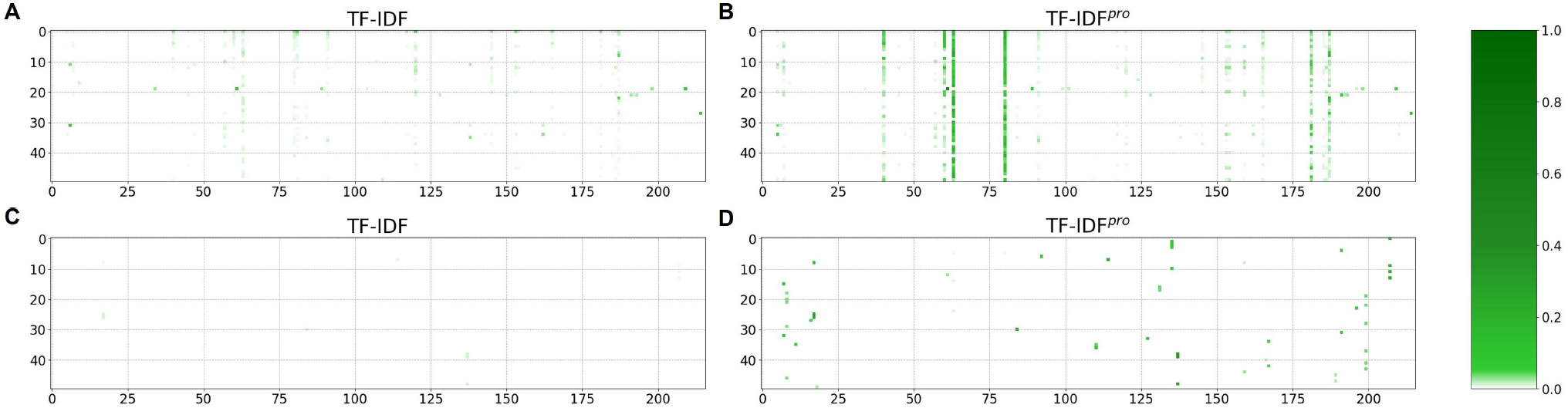
Comparison between the TF-IDF and TF-IDF^*pro*^ significance scores. (A) TF-IDF significance scores of the top 50 PCs, (B) TF-IDF^*pro*^ significance scores of the top 50 PCs, (C) TF-IDF significance scores of the last 50 PCs, and (D) TF-IDF^*pro*^ significance scores of the last 50 PCs. The PCs are sorted by the total number of protein sequences in each PC. In each subfigure, the x axis represents the index of each genus and the y axis represents the index of each PC.

Compared with TF-IDF, TF-IDF^*pro*^ can differentiate the significance scores of one PC to different genera more clearly. Especially among the PCs that contains smaller number of protein sequences, almost all the TF-IDF scores shown in Fig. 3(C) are zero because they appear less frequently than other PCs in each genus. On the contrary, TF-IDF^*pro*^ assigns higher significance scores for some PCs as their frequencies in one genus is higher than that in other genera. In summary, TF-IDF^*pro*^ is more suitable for our task of plasmid host range prediction.

Once we use the TF-IDF^*pro*^ to compute the importance of a PC to a genus, we use this information to assign possible host labels for a plasmid. With an input plasmid sequence *s*, we first identify the corresponding PCs of all encoded proteins in plasmid *s*. The set of all PCs is denoted as 𝒫. Then, the overall weight of all the PCs in 𝒫 to a genus *j*, i.e., 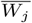 can be computed by averaging the TF-IDF^*pro*^ significance score of each PC to genus *j*. The equation is given below.

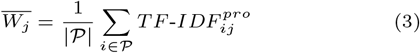

where | 𝒫 | is the total number of encoded protein sequences in the set 𝒫.

With the averaged weights, we compute the normalized weights 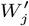 for all the genera as:

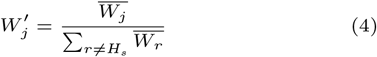

where *H*_*s*_ is is the genus-level host label of input plasmid *s*.

If the normalized weight of one genus is larger than a threshold, this genus is considered as extra pseudo label for plasmid sequence *s*. The assignment of pseudo labels is formulated as:

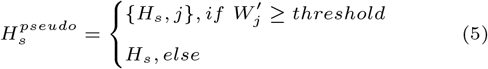

The default threshold is set to 0.5, indicating that each plasmid can have at most one pseudo label. By setting this stringent cutoff, we intend to ensure the high precision and reliability of generated pseudo label.

With the help of our pseudo label generation algorithm, we can provide additional label information for the training samples. In the experiment, we will evaluate the impact of our pseudo label generation algorithm to the performance of MOSTPLAS on plasmid host range prediction.

### Self-correction asymmetric loss

The other challenge of applying multi-label learning model on plasmid host range prediction is the significant imbalance between the number of positive and negative labels in the training samples. This inherent bias of the plasmid host labels leads to a result that the model parameter updating process pays more attention to the negative labels but overwhelms the contribution of rare positive labels. To cope with this problem, we design a self-correction asymmetric loss.

Traditional binary cross-entropy loss adopted for the training of multi-label learning model equally considers the contribution of positive labels and negative labels. Different from binary cross-entropy loss, our self-correction asymmetric loss takes advantage of the loss of positive labels to dominate model parameters updating process. Simultaneously, our self-correction asymmetric loss down-weights the contribution of easy samples during model training. Easy samples refer to the negative labels having predicted possibility closer to 0 and positive labels having predicted possibility closer to 1.

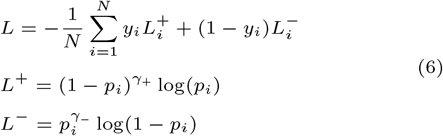

where *N* is the total number of genera, *y*_*i*_ is the binary label of input plasmid corresponding to genus *i, p*_*i*_ is the model prediction of input plasmid corresponding to genus *i*. As *p*_*i*_ ∈ [0, 1], we set *γ*_+_ *< γ*_−_ so that the weight of negative labels is smaller than that of positive labels.

In our sorted complete plasmid sequences, the number of negative labels is far more than that of positive labels, and the trained model is conserved to predict a large value of possibility that the input plasmid sequence belongs to one genus. Therefore, after adequate warming-up training epoch, if the model predicts a large possibility for a negative label, we consider it as a missing positive label. We design a self-correction mechanism to facilitate the training of multi-label learning model under the scenario that some of the host label of input plasmid sequence is possibly missing.

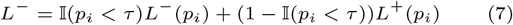

where *τ* is the threshold to identify a missing positive label, 𝕀(·) ∈ {0, 1} denotes the indicator function, if the condition holds, the value of the function is 1, otherwise the value is 0. We should notice that the self-correction mechanism is performed after several training epochs are finished, which we regard the model has certain discriminating ability at that moment.

In summary, our proposed self-correction asymmetric loss is formulated as:

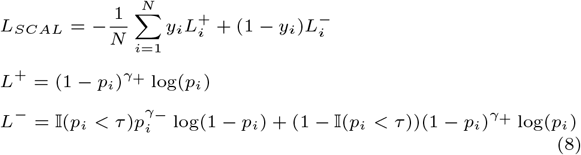

## Experiments

### Dataset

#### Multi-host plasmid test set

As the NCBI database only provides the bacterium where each plasmid sequence is isolated from, we cannot directly obtain the complete host labels set of each sequence. To screen possible plasmid sequences with multiple genus-level host labels from the complete plasmid sequences, we identify near-identical plasmid sequences with different genus-level host labels and combine their hosts as multi-host labels. Specifically, we employed BLASTN [29] to perform all-against-all alignments. In order to ensure higher reliability, we set a high threshold, where sequence pairs sharing alignments with higher than 99% identity and higher than 99% coverage of both query and matched sequences can be regarded as near identical. If the genus-level host labels of the two plasmid sequences are inconsistent, we consider these two sequences correspond to a cross-genus plasmid sequence with a minimal number of mutations. And thus, they share the combined genus-level host labels, which are the union set of their host labels.

By examining all BLASTN alignment results satisfying the 99% identity and coverage requirements, we obtained 1,391 plasmid sequences and their genus-level multi-host labels. We adopted all these sequences as the multi-host plasmid test set to evaluate the performance of MOSTPLAS, while the remainingplasmid sequences constitute the training set. The detailed host range information of each sequence in the multi-host plasmid test set is presented in the supplementary file.

#### Hi-C dataset

Hi-C sequencing can be used to capture three-dimensional genomic interactions between DNA molecules originating in the same cell, and thus is able to link plasmids to their bacterial hosts [30]. With metagenomic Hi-C data collected from wastewater, Stalder et al. successfully linked plasmid contigs to their host chromosomes and obtained 1307 groups of binned contigs [31]. In the same bin, the labels of all contigs are identical and annotated to the species level by Mash [32]. To identify plasmid contigs from all groups of binned contigs, we adopted MOB-recon [33] and Platon [34]. The intersection of the identified plasmids by the two tools were selected as the Hi-C dataset for our MOSTPLAS.

#### Mob-suite dataset

The authors of MOB-suite [33] performed a search of publications associated with plasmid accession numbers and manually identified and extracted the host range information of plasmid sequence referred in each article. As the host range of plasmid is experimentally determined and reported in literatures, we adopt this database to evaluate the performance of our model on real data. In our experiment, we removed the sequences with missing host range rank or host range. The detail host range information of the sorted 106 plasmid sequences is provided in the supplementary file.

#### DoriC dataset

DoriC was first launched in 2007 as a database of replication origins (Oris) in bacterial genomes [35]. In the latest version DoriC 12.0, the database includes Oris of plasmids. The Ori sequences in DoriC database are predicted based on Z-curve theory and comparative genomic method [36]. We employ this dataset to explore the biological characteristic of Oris in plasmids with multiple genus-level host labels.

### Evaluation metrics

To evaluate the performance of our proposed MOSTPLAS, we employ recall, precision, and F1-score as metrics. As the number of plasmids with different genus-level host labels is imbalanced, we adopted macro-averaging across the results of different genera and reported the mean value to represent the performance on the entire data. The formulations of macro-recall, macro-precision and F1-score are denoted as the following:

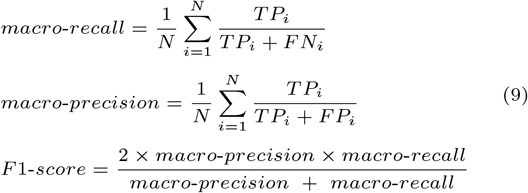

where *N* is the total number of genera, *TP*_*i*_ denotes correctly classified host label for genus *i, FN*_*i*_ is the number of missed host label, *FP*_*i*_ represents falsely predicted host label.

### Results on multi-host plasmid test set

#### Ablation study

We first performed ablation study to demonstrate the effectiveness of our proposed pseudo label generation algorithm and self-correction asymmetric loss. As mentioned in section 2.1, in our proposed MOSTPLAS, we designed three kinds of encoders. We employed all three encoders for this experiment, and the results are shown in Table 1.

**Table 1.**
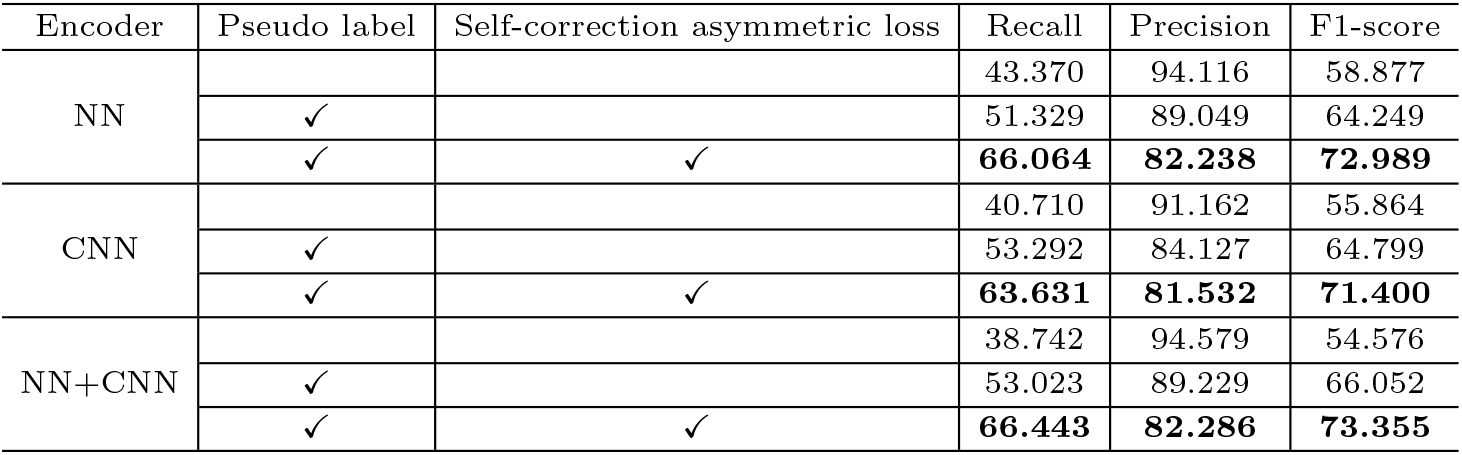
Ablation study results of our proposed pseudo label generation algorithm and self-correction asymmetric loss.

| Encoder | Pseudo label | Self-correction asymmetric loss | Recall | Precision | F1-score |
| --- | --- | --- | --- | --- | --- |
| NN |  |  | 43.370 | 94.116 | 58.877 |
|  | ✓ |  | 51.329 | 89.049 | 64.249 |
|  | ✓ | ✓ | **66.064** | **82.238** | **72.989** |
| CNN |  |  | 40.710 | 91.162 | 55.864 |
|  | ✓ |  | 53.292 | 84.127 | 64.799 |
|  | ✓ | ✓ | **63.631** | **81.532** | **71.400** |
| NN+CNN |  |  | 38.742 | 94.579 | 54.576 |
|  | ✓ |  | 53.023 | 89.229 | 66.052 |
|  | ✓ | ✓ | **66.443** | **82.286** | **73.355** |

After incorporating pseudo label, the recall of all three encoder models increased 7.959%, 12.582% and 14.281% respectively. Correspondingly, the improvements of F1-score were ranging from 5.372% to 11.476%. These results demonstrate the reliability of our proposed pseudo label generation algorithm.

In addition, if we replaced traditional binary cross-entropy loss with our self-correction asymmetric loss, all the three encoder models achieved the highest recall and F1-score. Among them, NN+CNN model performed the best, with 66.443% recall, 82.286% precision and 73.355% F1-score. This result shows that our proposed self-correction asymmetric loss helps multi-label learning model recognize more host labels of input plasmid sequences with a small sacrifice of precision.

According to Tabel 1, with the combination of pseudo label and self-correction asymmetric loss, all three encoders obtained a comparable performance on recall, precision and F1-score. The performance of our MOSTPLAS was robust to different options of encoders. In our proposed MOSTPLAS, we chose NN as the default encoder.

#### Comparison with alignment-based tool

We compared the performance of our models with BLAST, which is regarded as a benchmark of alignment-based tools. To obtain the results of BLAST, all the plasmid sequences in the multi-host plasmid test set are aligned with the plasmid sequences in the training set of our MOSTPLAS. The genus-level host label of the best matched sequences in the training set is regarded as the prediction of the query plasmid. The results are shown in Fig. 4.

**Fig. 4.**
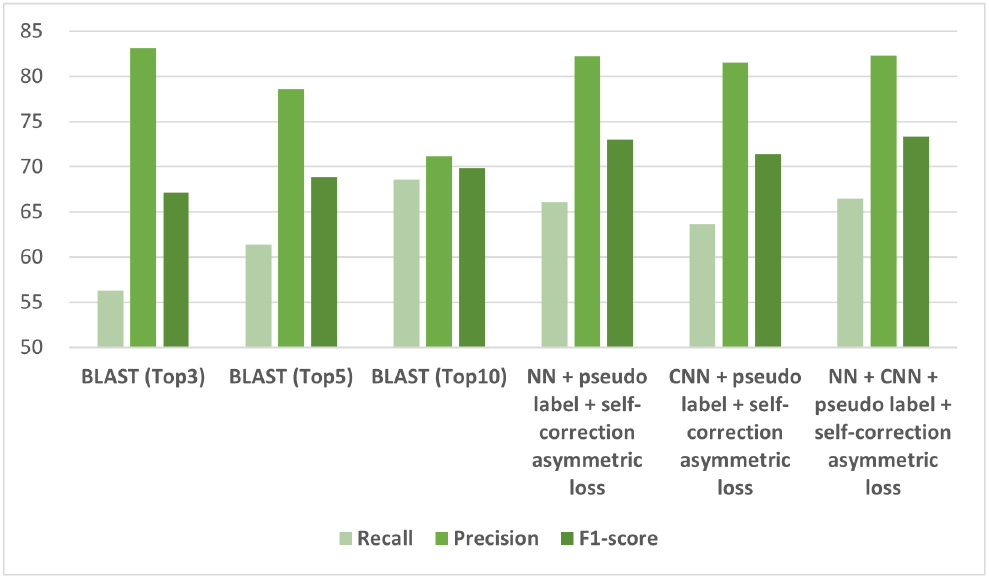
Performance comparison with alignment-based tool. BLAST (Top 3), BLAST (Top 5), and BLAST (Top 10) represent we chose the genus-level host labels of the top three, five, and ten best matched sequences as the predictions of BLAST.

Although our models obtain similar value of precision to BLAST (Top 3), the recall of our models is about 10% higher than that of BLAST. Similarly, when our models obtain similar value of recall to BLAST (Top 10), the precision of our models is also about 10% higher than that of BLAST. The lowest F1-score of our model (CNN) is still about 1.50% higher than the highest of BLAST (Top 10). These results demonstrate that our multi-label learning models have a better performance than BLAST on plasmid host range prediction.

#### Comparison with single-label learning tool

We also compared the performance of our MOSTPLAS with the single-label learning based tool HOTSPOT [13], which achieved the state-of-the-art performance on plasmid host range prediction. The results are shown in Fig. 5.

**Fig. 5.**
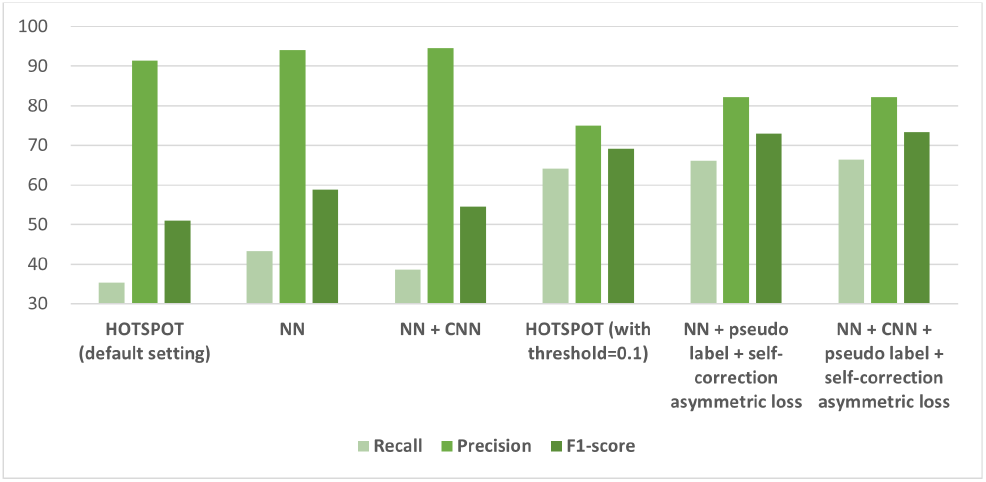
Performance comparison with single-label learning based tool. Beyond experiments with the default setting of HOTSPOT, we also evaluated the performance of HOTSPOT with a threshold of 0.1 to choose the predicted genus-level hosts.

The performance of HOTSPOT with default setting was comparable to our MOSTPLAS without incorporating pseudo label and self-correction asymmetric loss, as their recall were all lower than 50% and the F1-score were lower than 60%. If we modify the setting of HOTSPOT by setting a threshold of 0.1, which refers to taking all the genera with predicted probability larger than 0.1 as output, the recall increased significantly. However, the precision of HOTSPOT (with threshold=0.1) was about 8% lower and the F1-score was about 3% lower than our model. The results suggest that without the help of pseudo label generation algorithm and self-correction asymmetric loss, our MOSTPLAS perform fairly as single-label learning models. The combination of pseudo label generation algorithm and self-correction asymmetric loss plays an essential role in achieving better performance of multi-label learning model on plasmid host range prediction.

### Results on metagenomic data

To evaluate the performance of our MOSTPLAS on metagenomic data, we conducted experiments on the Hi-C dataset and compared the results with HOTSPOT. As the identified plasmid contigs in the Hi-C dataset were annotated with single-labels, we adopted accuracy as the evaluation metric. The accuracy is calculated as the number of correctly predicted plasmids divided by the total number of plasmids in the Hi-C dataset. The results are shown in Table 2.

**Table 2.**
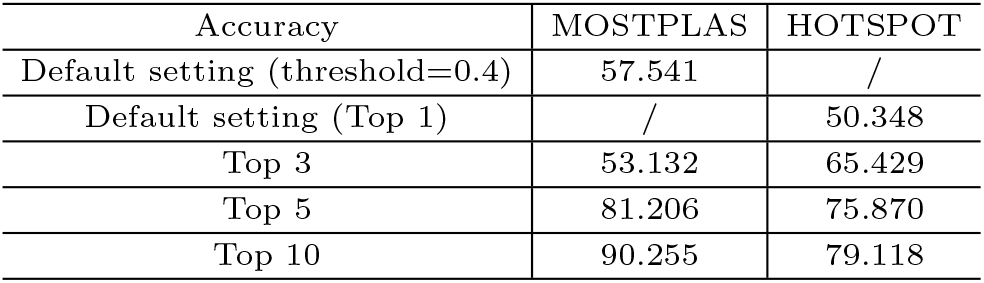
Performance comparison on the Hi-C dataset. In addition to experiments with the default setting of MOSTPLAS and HOTSPOT, we also compared the Top 3, Top 5 and Top 10 accuracy of MOSTPLAS and HOTSPOT.

| Accuracy | MOSTPLAS | HOTSPOT |
| --- | --- | --- |
| Default setting (threshold=0.4) | 57.541 | / |
| Default setting (Top 1) | / | 50.348 |
| Top 3 | 53.132 | 65.429 |
| Top 5 | 81.206 | 75.870 |
| Top 10 | 90.255 | 79.118 |

Under the default setting, our MOSTPLAS achieved a 7.193% higher accuracy than HOTSOPT. Although the Top3 classification accuracy of our MOSTPLAS was lower than that of HOTSPOT, the remaining Top 5 and Top 10 classification accuracy were much higher. The Top 10 accuracy of our MOSTPLAS was 90.255%, which indicated that we had high confidence that one of the top ten candidates was the host of input plasmid sequences. This result also demonstrated the effectiveness of our model on narrowing the query scope for plasmid with unknown host range in metagenomic data.

### Performance on plasmid sequences with experimentally determined host range

In the Mob-suite dataset, the host range label of each plasmid is annotated in different taxonomy levels, from superkingdom to family. However, our self-correction multi-label learning model performs genus-level plasmid host range prediction. To evaluate the performance of our model, for each predicted genus-level host label by our trained MOSTPLAS, we backtracked its corresponding taxonomy group that is of the same rank given by the dataset. In addition to recall and precision, we employ another metric, *exact match ratio* to evaluate the performance of our MOSTPLAS. Only if all the host labels of one plasmid sequence are predicted correctly, can we consider this plasmid sequence as a exactly matched sample. The prediction results are shown in Table 3.

**Table 3.** Results on real plasmid sequences with experimentally determined host range.

|  | Recall | Precision | Exact match ratio |
| --- | --- | --- | --- |
| Percentage | 107/115<br>(93.043) | 107/108<br>(99.074) | 101/106<br>(95.283) |

Among the 106 plasmid sequences in Mob-suite dataset, our models correctly predicted 101 samples, and the exactly matched ratio is above 95%. We further checked the samples with inconsistent prediction on their hosts, and presented their labels and predictions in Table 4.

**Table 4.**
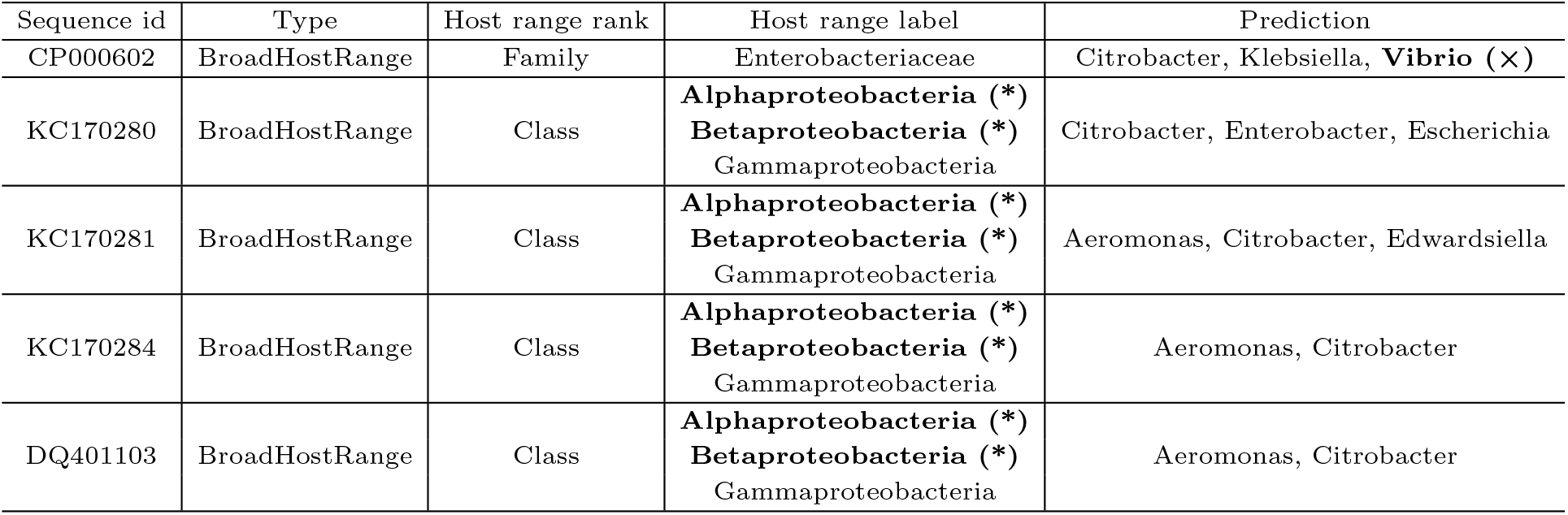
Details of the plasmid sequences without exactly matched prediction. (*) indicates a missed host label and (×) denotes a falsely predicted host label.

| Sequence id | Type | Host range rank | Host range label | Prediction |
| --- | --- | --- | --- | --- |
| CP000602 | BroadHostRange | Family | Enterobacteriaceae | Citrobacter, Klebsiella, **Vibrio (×)** |
| KC170280 | BroadHostRange | Class | **Alphaproteobacteria (\*)**<br>**Betaproteobacteria (\*)**<br>Gammaproteobacteria | Citrobacter, Enterobacter, Escherichia |
| KC170281 | BroadHostRange | Class | **Alphaproteobacteria (\*)**<br>**Betaproteobacteria (\*)**<br>Gammaproteobacteria | Aeromonas, Citrobacter, Edwardsiella |
| KC170284 | BroadHostRange | Class | **Alphaproteobacteria (\*)**<br>**Betaproteobacteria (\*)**<br>Gammaproteobacteria | Aeromonas, Citrobacter |
| DQ401103 | BroadHostRange | Class | **Alphaproteobacteria (\*)**<br>**Betaproteobacteria (\*)**<br>Gammaproteobacteria | Aeromonas, Citrobacter |

According to Table 4, our model missed two out of three host labels of four plasmid sequences with class level annotations. In addition, MOSTPLAS made a false prediction for plasmid CP000602. Our model predicted three genus-level host labels for this plasmid sequences. Among them, *Citrobacter* and *Klebsiella* are two genera belonging to *Enterobacteriaceae*, which is the family level label provided by the Mob-suite dataset. Besides, our model predicted another genus label *Vibrio*, which belongs to another family level label *Vibrionaceae*, but they share the same ancestor in the class level, as shown in Fig. 6.

**Fig. 6.**
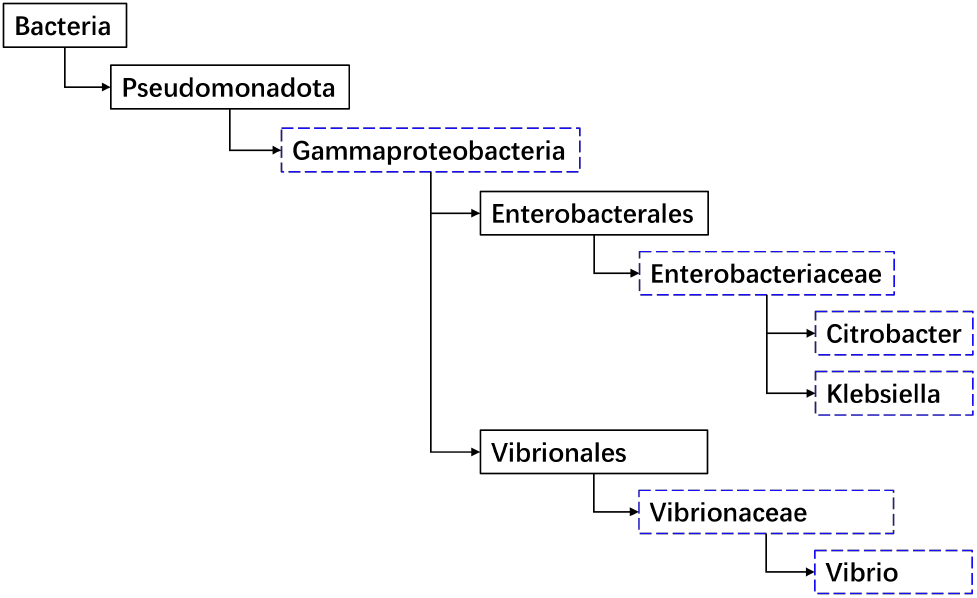
Phylogenetic tree of the predicted genus-level plasmid hosts *Citrobacter, Klebsiella*, and *Vibrio*.

### Exploration on the DoriC dataset

Previous research concludes that BHR plasmids may contain multiple basic replicons and different replicons will be activated in different hosts, which helps extend the plasmid host range [37]. To examine this characteristic using the identified multi-host plasmids identified by our tool, we utilized DoriC12.0 [35], a database that collects several Ori sequences from plasmids. In this experiment, we initially use our trained MOSTPLAS to predict the genus-level host labels for all complete plasmid sequences downloaded from the NCBI RefSeq database. Subsequently, we selected the sequences with at least 2 host labels and performed alignments using BLASTN against the Ori sequences in the DoriC12.0 database. For each alignment of a plasmid sequence, if both the identity and coverage exceeded 95%, we consider this plasmid includes the matched Ori sequence. Overall, we obtain 19,241 sequences with at least 2 host labels, and the distribution of their Ori numbers is illustrated in Fig. 7.

**Fig. 7.**
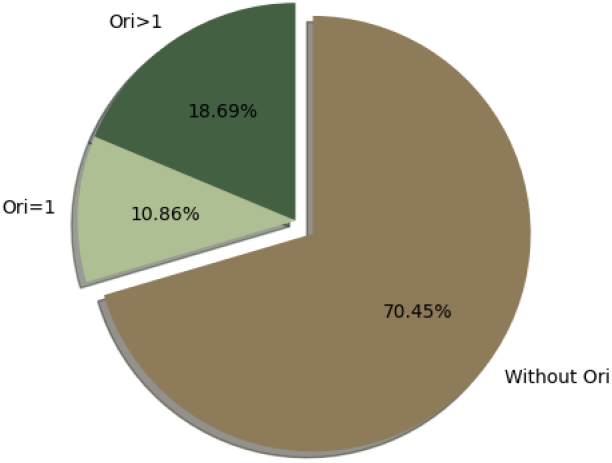
The distribution of the Ori number in the predicted plasmids with more than two genus-level host labels.

Approximately 70.5% plasmids in our experiment had no alignments with the OriC sequences present in the DoriC12.0 database. This discrepancy can be attributed to the limited coverage of the database, which currently includes only 1,184 Ori sequences extracted from plasmids. Considering the rapid increase in the number of complete plasmid sequences available in the RefSeq database, it is likely that numerous Ori sequences in plasmids remain undiscovered. Additionally, the experiment results reveal that around 7.8% more plasmids contain multiple Ori sequences compared to those with a single Ori. This observation suggests that plasmid with multiple genus-level host label tends to have more replicons, which is consistent with existing literature [37]. This experiment underscores the importance of identifying Ori sequences within plasmids.

### Study of biological characteristic of plasmids with multiple genus-level host labels

Several reviews on plasmid biology [25, 38] suggest that transmissible plasmids often carry genes that facilitate their establishments in newly infected cells, especially when the cells differ genetically from the previous host. Building upon this insight, we consider to investigate whether the host range of a longer plasmid sequence is wider than or the same as that of a shorter plasmid, which serves as its subsequence. To accomplish this, we utilized all the complete plasmid sequences extracted from the NCBI RefSeq database and performed all-against-all BLASTN alignments between them. If the alignments between two plasmid sequences fulfilled two criteria: 1) the identity was greater than 99% and 2) the coverage of short sequence exceeded 99%, we considered the shorter sequence as the subsequence of the longer one. The host range of all complete plasmid sequences was predicted using our trained MOSTPLAS model. We sorted the host range relationship between all pairs of plasmid sequences, and the results were presented in Fig. 8. In Fig. 8(A)-(C), a significant portion of the plasmid sequence pairs exhibit the same host range, with the percentage exceeding 40%. When the coverage of the subsequence within the long sequence is below 75% or 50%, approximately 80% plasmid sequence pairs align with our assumption that their host ranges have an inclusion or consistent relation. However, if the coverage of the subsequence within the long sequence dropped below 25%, only around 60% plasmid sequence pairs satisfied our assumption.

**Fig. 8.**
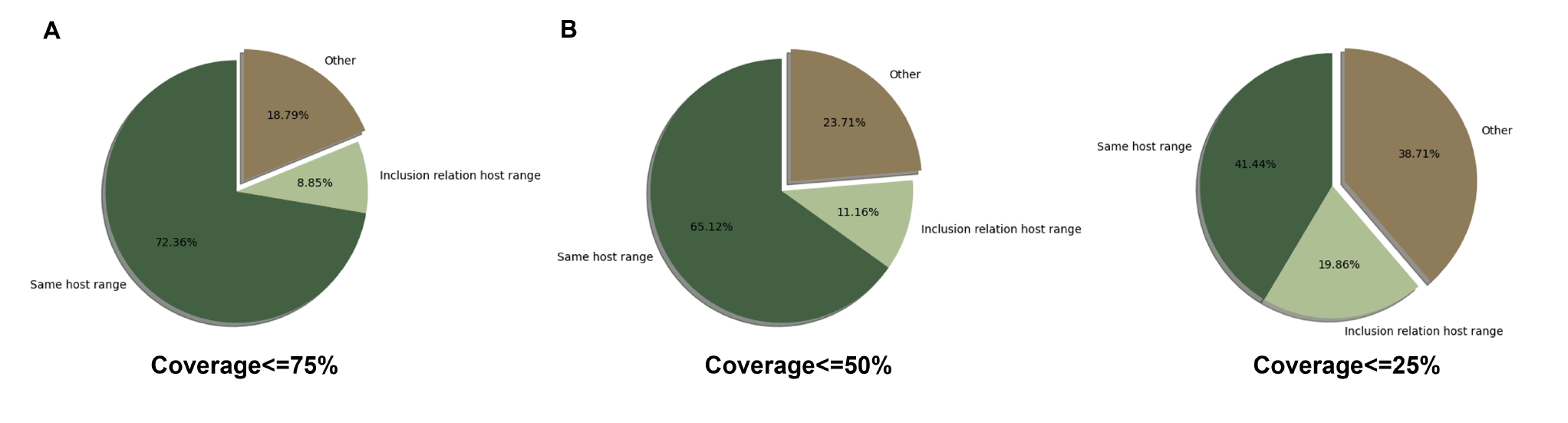
Host label comparison results of plasmid sequence pairs. (A) The coverage of the subsequence in the long sequences is ≤ 75%, (B) The coverage of the subsequence in the long sequences is ≤ 50%, and (C) The coverage of the subsequence in the long sequences is ≤ 25%. We chose plasmid sequence pairs where short sequence is the subsequence of the long sequence. Same host range group represented the host range of the long sequence and its subsequence were identical. If the host range of the subsequence was the subset of the host range of the long sequence, we considered their host ranges had an inclusion relation. If the host range of the subsequence did not satisfy the previous two conditions, this plasmid sequence pair was categorized into the other group.

## Conclusion and Discussion

In this paper, we present a novel self-correction multi-label learning model called MOSTPLAS for genus-level plasmid host range prediction. To the best of our knowledge, this is the first attempt to apply a multi-label learning model to this task. In MOSTPLAS, we design a pseudo label learning algorithms to mitigate the limitation of no available database providing complete host range of BHR plasmid sequences for deep learning models to extract discriminative feature representation for each genus. We also introduce a self-correction asymmetric loss that adjust the preference of traditional binary cross-entropy loss on the negative labels during model parameters updating. The experiment results on multi-host plasmid test set generated from the NCBI RefSeq database, metagenomic data, and real-world plasmid sequences with experimentally determined host range demonstrate the effectiveness of our MOSTPLAS.

Although MOSTPLAS showed promising performance under the challenging scenario that the host labels of plasmid sequences include unknown false negative labels, there are still areas for improvement in future research. Firstly, MOSTPLAS requires complete plasmid sequences as input, as there is currently insufficient evidence showing the host range relationship between complete BHR plasmid sequences and their corresponding plasmid contigs. Moreover, the host labels of the plasmid sequences in our multi-host plasmid test set were determined with a strict BLASTN alignment threshold. The other predicted non-host bacteria still contain some potential false negative labels. In future work, it would be valuable to investigate the mechanism how the origins of replication, transposons as well as other mobile genes contribute to the host adaption of plasmids function, which may provide us with more clues on determining the host range of plasmid contigs.

## Supporting information

Supplementary_1_multi-host plasmid test set

Supplementary_2_mob_suit_sorted

## Data availability

MOSTPLAS is implemented with Python, which can be downloaded at https://github.com/wzou96/MOSTPLAS.

## Supplementary data

Supplementary Data are available at NAR Online.

## Acknowledgments

The authors thank the anonymous reviewers for their valuable suggestions. This work is supported by City University of Hong Kong; Hong Kong Innovation and Technology Commission (InnoHK Project CIMDA).

