## Supplementary_3_supplementary file for "MOSTPLAS: A Self-correction Multi-label Learning Model for Plasmid Host Range Prediction"

### 1. Model architecture

In our MOSTPLAS, we designed three different kinds of encoder for feature extraction: Neural Network (NN), Convolutional Neural Network (CNN) and a multi-model integration framework (NN+CNN). These encoders were then cascaded with a fully-connect (FC) layer and a Sigmoid activation layer together to formed an end-to-end plasmid host range prediction model.

In bioinformatic, k-mer frequency is a widely used feature for feature analysis [1,2]. For NN encoder, we used the 4-mer reverse complementary frequency vector of input plasmid sequence as model input. The dimension of input vector is 136. The encoder is composed of 4 FC layers and the hidden neuron number is set as 2048. 1024, 512 and 256 respectively for each layer. Among two FC layers, we incorporated a BatchNorm1d layer and a ReLU activation layer.

For CNN encoder, we first used Prodigal to predict the start and end site of all encoded genes from input plasmid sequences. Then, we converted the DNA sequences of all encoded gene into 4-mer reverse complementary frequency vectors. At last, we concatenated all the vectors into one matrix according to gene arrangement order and employ the matrix as model input. We restricted the maximum length of the matrix to 200. For plasmid sequences with less than 200 encode genes, we use zero-padding to make sure the input matrix has uniform size as 200×136.

Inspired by the CNN model adopted for text classification [3], the encoder in our CNN variant model includes four convolutional layers and each convolutional layer is cascaded with a BatchNorm2d layer and a ReLU activation layer. As the encoder performs convolution operation among contiguous genes for feature extraction, the kernel size of each convolutional layer is set as 2×136, 3×136, 4×136 and 5×136, respectively. The output channel of all convolutional layers is set as 256 and the stride is set to 1. The feature maps output from all convolutional layers are then passed to a global max pooling layer and converted into a vector with dimension as 256. The adopted four vectors are further concatenated together and sent to a FC layer with 256 hidden neurons.

For NN+CNN encoder, we aim to leverage the strengths of both NN and CNN models. The integration of different models can be achieved through feature-level fusion [4] and decision-level fusion [5]. Based on prior research [6], decision-level fusion has demonstrated superior performance compared to feature-level fusion strategies. Therefore, in our NN+CNN variant model, we perform decision-level fusion to integrate the output of NN and CNN models, which is formulated as:

$$p_i = \lambda p_i^{NN} + (1-\lambda) p_i^{CNN}$$
$$L = \lambda L^{NN} + (1-\lambda) L^{CNN}$$

where $p_i^{NN}$ and $p_i^{CNN}$ are the output of NN model and CNN model, $L^{NN}$ and

$L^{CNN}$ are the loss of NN model and CNN model, $\lambda$ is a hyper-parameter to balance the contribution of two models. In this work, we set $\lambda$ as 0.6.

### 2. Training setting

Our self-correction multi-label learning model is implemented with Python 3.10.13 and Pytorch 1.12.1 deep learning platform. We also adopt one NVIDIA GeForce RTX 3090 GPU to speed up model training. We use Adam as the optimizer and the initial learning rate is set as 1e-3. The learning rate is decreased to 1/2 of initial learning rate after 50 training epochs and the entire training epoch is set as 100. In our proposed Self-correction asymmetric loss, the hyperparameters $\gamma_+$ and $\gamma_-$ that adopt to weight the contribution of positive labels and negative labels are set as 1 and 2 respectively. The label correction is performed after 15 epochs and the threshold $\tau$ to identify a missing positive label is set as 0.7.

### 3. Evaluation of the reliability of pseudo label generation algorithm

We employed the multi-host plasmid test set to explore the reliability of our pseudo label generation algorithm. For each plasmid sequence in the dataset, we removed the genus level host labels provided by NCBI RefSeq database and evaluated the performance on the remaining labels.

Compared with TF-IDF$^{pro}$, the pseudo label generation procedure of TF-IDF was determined by the averaged significance of all protein sequences of one plasmid. For all the genera, if the significance of one genus is larger than the significance of the host label of the plasmid, this genus is chosen as the pseudo label. The results are shown in Table 1.

Table 1: Performance comparison between different pseudo label generation algorithms.

| Method | Percentage of retrieved BLAST label in pseudo label | Percentage of pseudo label consistent with BLAST label |
|---|---|---|
| TF-IDF | 36.857 | 25.410 |
| TF-IDF (normalization) | 17.701 | 54.235 |
| **TF-IDF$^{pro}$** | **21.048** | **83.301** |

Compared with TF-IDF algorithm, the retrieved BLAST label in the pseudo label generated by TF-IDF$^{pro}$ measurement decreased about 15.8%. However, the consistency between the pseudo label generated by TF-IDF$^{pro}$ measurement and BLAST label improved about 57.9% and reached more than 83%. We also performed normalization on TF-IDF scores and generated the pseudo labels following TF-IDF$^{pro}$ procedures. When we incorporated TF-IDF scores with normalization, the percentage of retrieved labels decreased about 19% and the consistency with BLAST labels improved 29%, which demonstrated that our pseudo labels generation pipeline ensured a high precision.

Since the pseudo labels are adopted as extra supervision on model training, they should be exact enough or would introduce noise to the training samples. A large amount of falsely labelled samples may significantly degrade model performance. To this end, we set a strict threshold to ensure the high precision of our generated pseudo labels. The high precision demonstrates our protein TF-IDF algorithm is available to generate credible pseudo labels and help boost the performance of multi-label learning models on plasmid host prediction task. We also present some examples of the generated pseudo labels in Table 2.

Table 2: Examples of pseudo labels generated by TF-IDF and our TF-IDF$^{pro}$. × denotes falsely predicted labels.

| Sequence id | Isolated organism | BLAST label | Pseudo label generated by TF-IDF | Pseudo label generated by TF-IDF$^{pro}$ |
|---|---|---|---|---|
| NZ_AP026682 | Salmonella | Enterobacter<br>Leclercia<br>Salmonella | Enterobacter<br>Leclercia<br>Phytobacter **(×)**<br>Salmonella **(×)**<br>Serratia | Enterobacter<br>Salmonella |
| NZ_OW967522 | Klebsiella | Enterobacter<br>Klebsiella<br>Phytobacter | Citrobacter **(×)**<br>Enterobacter<br>Klebsiella<br>Leclercia **(×)**<br>Phytobacter<br>Salmonella **(×)**<br>Serratia **(×)** | Enterobacter<br>Klebsiella |

In Table 2, although TF-IDF successfully recognized all the genus level host labels of the two plasmid sequences, it also made a large number of predictions that are not included in the BLAST label set. TF-IDF$^{pro}$ algorithm tend to make fewer predictions but ensure the high precision of the pseudo labels. This comparison result demonstrated the reliability of the pseudo labels generated by TF-IDF$^{pro}$ significance scores.

[2] H. Yan, A. Bombarely and S. Li, “DeepTE: a computational method for de novo classification of transposons with convolutional neural network,” Bioinformatics,

2020, 36(15): 4269-4275.

[3] S. Wang, M. Huang and Z. Deng, “Densely connected CNN with multi-scale feature attention for text classification,” International Joint Conference on Artificial Intelligence (IJCAI). 2018, 18: 4468-4474.

[4] H. D. Nguyen, S. H. Kim, G. S. Lee, et al., “Facial expression recognition using a temporal ensemble of multi-level convolutional neural networks,” IEEE Transactions on Affective Computing, 2019, 13(1): 226-237.

[5] H. Jung, S. Lee, J. Yim, et al., “Joint fine-tuning in deep neural networks for facial expression recognition,” Proceedings of the IEEE international conference on computer vision (ICCV), 2015: 2983-2991.

[6] Z. Zhao, Q. Liu and S. Wang, “Learning deep global multi-scale and local attention features for facial expression recognition in the wild,” IEEE Transactions on Image Processing, 2021, 30: 6544-6556.
